# Rhythmic modulation of visual perception by continuous rhythmic auditory stimulation

**DOI:** 10.1101/2020.10.30.362467

**Authors:** Anna-Katharina R. Bauer, Freek van Ede, Andrew J. Quinn, Anna C. Nobre

## Abstract

At any given moment our sensory systems receive multiple, often rhythmic, inputs from the environment. Processing of temporally structured events in one sensory modality can guide both behavioural and neural processing of events in other sensory modalities, but whether this occurs remains unclear. Here, we used human electroencephalography (EEG) to test the cross-modal influences of a continuous auditory frequency-modulated (FM) sound on visual perception and visual cortical activity. We report systematic fluctuations in perceptual discrimination of brief visual stimuli in line with the phase of the FM sound. We further show that this rhythmic modulation in visual perception is related to an accompanying rhythmic modulation of neural activity recorded over visual areas. Importantly, in our task, perceptual and neural visual modulations occurred without any abrupt and salient onsets in the energy of the auditory stimulation and without any rhythmic structure in the visual stimulus. As such, the results provide a critical validation for the existence and functional role of cross-modal entrainment and demonstrates its utility for organising the perception of multisensory stimulation in the natural environment.

**Significance Statement:** Our sensory environment is filled with rhythmic structures that are often multi-sensory in nature. Here we show that the alignment of neural activity to the phase of an auditory frequency-modulated sound has cross-modal consequences for vision: yielding systematic fluctuations in perceptual discrimination of brief visual stimuli that is mediated by accompanying rhythmic modulation of neural activity recorded over visual areas. These cross-modal effects on visual neural activity and perception occurred without any abrupt and salient onsets in the energy of the auditory stimulation and without any rhythmic structure in the visual stimulus. The current work shows that continuous auditory fluctuations in the natural environment can provide a pacing signal for neural activity and perception across the senses.

## Introduction

Our sensory environment is filled with rhythmic structure to which neural activity can become synchronised. This synchronisation of neural activity and rhythmic structures in the environment is often referred to as “neural entrainment”, a process by which two self-sustained oscillations become coupled via phase and/or frequency adjustment (Pikovsky et al., 2003; Lakatos et al., 2008; Thut et al., 2011). Low-frequency neural entrainment has been suggested as an important mechanism for enabling cross-modal influences by facilitating the transfer of information across sensory modalities (Van Atteveldt et al., 2014; Simon and Wallace, 2017; Keil and Senkowski, 2018; Lakatos et al., 2019; Bauer et al., 2020).

To date, studies that have investigated cross-modal influences of auditory rhythms on visual perception have relied on rhythmic auditory streams with identifiable onsets and offsets, such as individual transient events (tones) or amplitude modulations (Lakatos et al., 2008; Bolger et al., 2013; Brochard et al., 2013; Miller et al., 2013; Escoffier et al., 2015; Simon and Wallace, 2017; Barnhart et al., 2018; Bauer et al., 2020; Chow et al., 2020). However, such onsets may cause cross-modal influences simply because they are salient or because the individual events that make up such rhythms may repeatedly evoke cross-modal phase-resets of neural activity (Fiebelkorn et al., 2011, 2013; Naue et al., 2011; Romei et al., 2012; Mercier et al., 2013; Diederich et al., 2014; Cecere et al., 2015; Keil and Senkowski, 2017; Mégevand et al., 2020). Thus, any observed behavioural or neural modulations may not directly or only partly entail true “entrainment” of neural activity (Breska and Deouell, 2017; Haegens and Zion Golumbic, 2017; Novembre and Iannetti, 2018; Doelling et al., 2019; Helfrich et al., 2019; Obleser and Kayser, 2019). To bypass this complication when interpreting cross-modal influences of auditory rhythmic stimulation on visual performance and neural activity, we used a continuous auditory stimulus for which periodicity was conveyed via frequency-modulation (FM). The advantages of using an FM-sound is that this type of auditory stimulation conveys clear “rhythmicity” in the perception of the listener, while keeping the overall “energy” (amplitude) of the sound constant over time (as shown in Figure 1A). Critically, the FM tone allows the to-be-discriminated targets to be placed at any phase along the auditory stimulation without the interference from perceptually abrupt discontinuous events.

**Figure 1.**
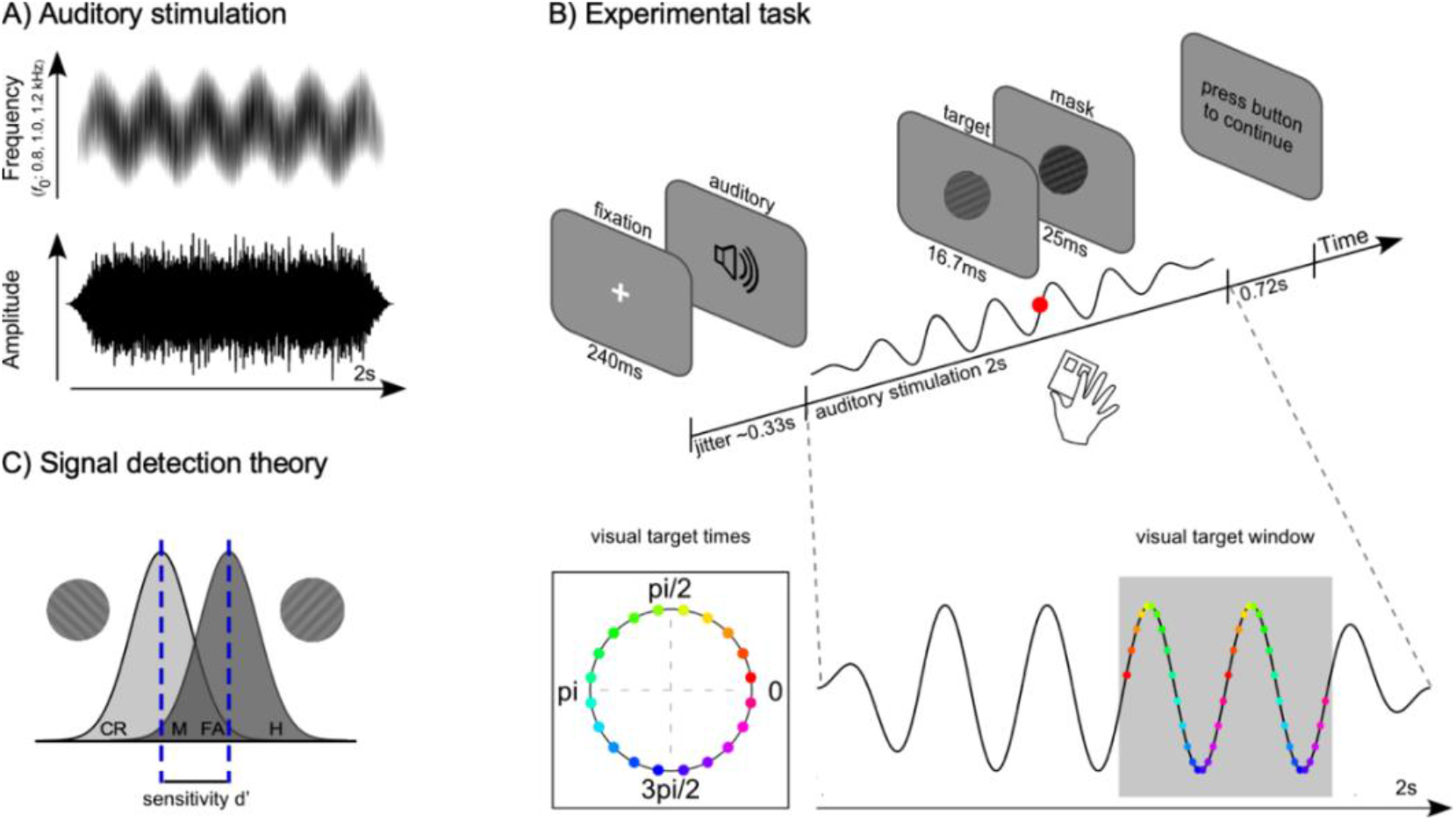
FM-sound characteristics, experimental task, and illustration of signal detection theory. A) Auditory stimulation. Driving stimulus characteristics of the 3-Hz frequency-modulated (FM) stimulation. Periodicity was conveyed by fluctuations in frequency, but without fluctuations in amplitude (or “stimulus energy”). The centre frequency *f*_0_ was randomized from trial to trial and could take on one of three values: 800 Hz, 1000 Hz, and 1200 Hz. B) Experimental task. Schematic of a single trial. Each trial started with a fixation cross after which the auditory stimulation started. After an interval of at least 1000 ms, a single Gabor grating, oriented either 45- or 135-degrees, appeared briefly at a personalised contrast (16.7 ms) and was immediately masked (25 ms). Participants indicated the orientation of the grating via a button press. The key manipulation was that Gabor gratings were systematically presented with respect to the phase angle of the 3-Hz FM stimulation (20 visual target times in total; first visual target time set to 8°; distance between visual targets 18°). The initial phase of the auditory stimulus was varied on a trial by trial basis and could take one of four values: 0, pi/2, pi, and 3pi/2. Gabor gratings were presented after 1 s of auditory stimulation and could occur across two cycles. C) Signal detection theory. For each visual target time, we computed visual target sensitivity (d’) based on the hit rate (H) from the 45-degree orientation condition and the false-alarm rate (FA) from the 135-degree orientation condition (CR: correct rejection; M: miss).

In previous uni-modal work, we and others have demonstrated that a continuous auditory FM stimulation can profoundly impact the ability to detect near-threshold auditory targets embedded in the auditory stream, such that detection performance varies systematically with the phase of the FM-sound (Henry and Obleser, 2012; Henry et al., 2014, 2017; Bauer et al., 2018). Further, previous studies investigating cross-modal auditory-to-visual influences focused either on behavioural (Miller et al., 2013; Barnhart et al., 2018) or neural activity (Lakatos et al., 2008; Escoffier et al., 2015). Here, we looked at both, asking whether a continuous FM-sound can influence visual perception, and whether this influence could be accounted for by the concomitant cross-modal influence on visual brain activity.”

To test for cross-modal auditory-to-visual influences, participants (*N* = 28) identified the orientation of a briefly presented visual Gabor grating, either rotated 45- or 135-degree, embedded within a two-second 3-Hz frequency-modulated (FM) stimulus (Figure 1A; <u>Sound</u>) while electroencephalography (EEG) was recorded. Critically, visual targets were presented at different times, relative to the phase of the continuous FM auditory stimulation (see Figure 1B: visual target times; <u>Video</u>). We predicted that ongoing neural oscillations in auditory cortices would entrain to the 3-Hz FM stimulation. Building on predictions from unimodal studies (Henry and Obleser, 2012; Henry et al., 2014, 2017; Bauer et al., 2018), we were here specifically interested in the cross-modal influences of the continuous 3-Hz FM auditory stimulation. We tested whether the perceptually varying auditory stimulus would lead to rhythmic modulation of visual perception, and sought evidence for rhythmic entrainment of neural activity in visual brain areas that could account for such rhythmic modulation of visual perception.

## Materials and Methods

### Participants

The study was approved by the Central University Research Ethics Committee of the University of Oxford and was conducted in accordance with the Declaration of Helsinki. Thirty healthy human volunteers (18 female, 12 male) participated in the study after providing written informed consent. Sample size was set a-priori based on our experience with similar tasks (Bauer et al., 2018). Data from two out of the 30 participants were discarded. One participant did not follow task instructions and pressed response buttons several times, and at random points, throughout many of the trials. The behavioural performance of the other participant was below chance (negative d’ values) indicating response confusion or misunderstanding of the task. Analyses are based on the 28 remaining participants (18 female; age range = 20 to 35, *M*_age_ = 27.4, *SD* = 4.0). One participant was left-handed and nine participants were ambidextrous as indicated by the Edinburgh Handedness Inventory (EHI; Oldfield, 1971). All participants had normal or corrected-to-normal vision and reported no history of hearing, neurological, or psychiatric disorders. Participants received financial compensation of £15 per hour.

### Stimuli

Auditory stimuli were generated using MATLAB software (Mathworks, Inc.). Frequency-modulated (FM) auditory stimuli were two-second narrow-band noises modulated at a rate of 3 Hz with a modulation depth of 37.5% and sampled at 48 kHz (similar to previous studies: Henry and Obleser, 2012; Henry et al., 2017; Bauer et al., 2018; <u>Sound</u>). Periodicity was conveyed via fluctuations in frequency (pitch) instead of amplitude fluctuations (see Figure 1A). Critically, this allowed us to induce a 3-Hz rhythm without changing the total energy of the auditory stimulation over time. The FM signal was faded in and out by using a 333-ms Hanning ramp, corresponding to one cycle of the 3-Hz FM stimulation. The centre frequency of the complex carrier signal was randomised from trial to trial and could take one of three values (800 Hz, 1000 Hz, 1200 Hz). The carrier signals were centred on one of these three frequencies and were constructed by adding 30 frequency components sampled from a uniform distribution with a 500-Hz range. The onset phase of the stimulus varied on a trial-by-trial basis and could take on of four values (0, pi/2, pi, and 3pi/2). All stimuli were normalised with respect to the root-mean square. Auditory stimuli were presented binaurally over EARTone 3A insert earphones (3M Auditory Systems, Indianapolis, United States) at a comfortable listening level (self-adjusted by each listener).

Visual target stimuli were generated using R software (version 1.2.1335, R Core Team, 2016, Vienna, Austria). They consisted of Gabor gratings with a spatial frequency of 2.5 cycles per degree of visual angle and tilted at either 45 or 135 degrees (pi/4 and 3pi/4 respectively). Gratings had a diameter of 2 degrees visual angle and were presented foveally (screen resolution: 1920 x 1080; monitor refresh rate: 120 Hz). The masking stimulus was generated by overlaying two Gabor gratings with the same orientation as the target stimuli, either 45 or 135 degree, but with a higher spatial frequency of 10 cycles per degree of visual angle. In addition, the mask was convolved with a white Gaussian kernel. The Gaussian kernel was generated in R using the formula provided by the Visual Stimulus Generation Toolkit (Neurobehavioral Systems Inc., Albany, US):

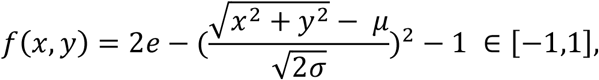

where the coordinates x = 0 and y = 0 correspond to the centre of the screen as well as the centre of the gaussian kernel. The parameter *μ* was constant and set to 0 and the parameter *σ* which controlled the size of the radius was set to 0.30 corresponding to a visual angle of 2 degree at a screen resolution of 1920×1080. Targets and masks were presented on a grey background (RGB: 109, 109, 109).

Critically, we manipulated when the visual targets were presented along the phase of the auditory 3-Hz FM stimulation. In particular, Gabor gratings occurred in one of 20 equally spaced times that were defined relative to the phase of the 3-Hz FM cycle of the auditory stimulation (Figure 1B: visual target times). The first phase in which a visual target could appear relative to the auditory stimulation was 8 degrees into the 3-Hz FM cycle, and subsequent targets occurred in steps of 18 degrees around the cycle. Gabor gratings were presented after a minimum of one second of the 3-Hz FM stimulation, and potential visual targets were distributed across two cycles of the auditory stimulation to reduce predictability of its exact temporal onset within the auditory stream (Figure 1B: visual target window: 1000 to 1666 ms).

To ensure accurate alignment between visual targets and the 20 possible phases of the auditory stimulus, Gabor gratings were embedded within videos with a duration of 2 seconds (240 frames), corresponding to the length of the auditory stimulation. Targets were presented for 16.7 ms (2 frames) and were immediately followed by the mask, which was presented for 25 ms (3 frames). While the contrast of the Gabor grating was individually adjusted for each participant (see Calibration procedure), the contrast of the mask was constant across participants. All other images were blank grey frames (RGB: 109,109,109). Single frames were exported to FFmpeg (http://ffmpeg.org) and converted into videos (video codec: libxvid; <u>Video</u> containing both the auditory stimulation and the visual target).

A second set of videos with a duration of 1 second (120 frames) were created for establishing a participant’s individual Gabor contrast (see section Calibration procedure). Gabor gratings were inserted after 500 ms for 16.7ms (2 frames) and followed by mask presented for 25 ms (3 frames). The remaining frames were blank grey frames (RGB: 109, 109, 109). Single frames were subsequently exported to FFmpeg and converted into videos.

### Procedure

Each session started with a calibration session to estimate the Gabor contrast threshold for each participant and was followed by a practice block before the main task started. Participants sat in a dimly lit and electrically shielded sound-attenuated booth, approximately 95 cm in front of the screen. Behavioural data were recorded online by Presentation software (version 18.3.06.02.16, Neurobehavioral Systems Inc., Albany, US). Responses were collected on a standard keyboard.

### Calibration procedure

Threshold contrasts for perceiving the Gabor gratings were first titrated for each participant using a two-alternative forced choice task (2AFC). A three-down one-up rule was implemented aiming for an accuracy of ~70% (Levitt, 1971). In each trial, a Gabor grating was presented for 16.7 ms, tilted 45 or 135 degree, followed immediately by a mask of 25 ms duration. Participants had to indicate the orientation of the Gabor grating by pressing the right (45 degree) or left (135 degree) arrow key on the keyboard. Each trial started with a variable delay centred on 333 ms, which was followed by a white fixation cross that was presented for 240 ms. The video, containing the Gabor grating and mask, started after another variable interval (jitter centred on 333 ms) and lasted for 1000 ms. Participants were then prompted with a response screen and the next trial started after participants indicated the orientation of the Gabor grating. Participants completed three blocks of the 2AFC task, lasting about five minutes each. In each block twelve reversals were completed and the threshold contrast was determined by the average threshold of the last eight reversals (procedure similar to: Henry and Obleser, 2012; Bauer et al., 2018). The final threshold contrast for the Gabor gratings was defined as the arithmetic average of the individual estimates for each of the three blocks.

### Cross-modal task

In the main task (Figure 1B for a schematic), participants had to discriminate the orientation of the Gabor gratings (45 or 135 degree). Participants were asked to listen to the auditory stimulation and to indicate the orientation of the Gabor grating as quickly as possible by pressing the right or left arrow key on the keyboard, corresponding to the 45- and 135-degree orientation respectively. Visual targets were presented for 16.7 ms at the individually established contrast and immediately masked for 25 ms. In each trial, only one visual target was presented.

The cross-modal task was self-paced, in that participants could initiate each trial on their own by pressing the space bar. After a variable delay, centred on 333 ms (1 cycle of the 3-Hz FM stimulation), a white fixation cross appeared for 240 ms. The fixation cross was followed by another variable interval centred on 333 ms after which the auditory-visual stimulation started. The auditory stimulation and the video, containing the Gabor grating and mask, started simultaneously and lasted for 2000 ms. The auditory-visual stimulation was followed by a 720 ms blank screen in which potential responses would still be counted. The participants were then prompted on the screen to initiate the next trial by pressing the space bar. Participants were explicitly instructed to maintain fixation on the middle of the screen after the fixation cross disappeared and were asked to proceed in a timely manner from trial to trial.

Participants completed a short practice block (12 trials) in which they received feedback on the screen regarding the correct identification of the Gabor orientation. No feedback was presented during the main task. In the main task, each participant completed 640 trials, resulting in 32 trials for each visual target time (16 per Gabor orientation) across the two cycles. The main task was presented in six blocks of ~10 min each with breaks of self-determined length in-between blocks and lasted on average 60 minutes (*SD* = 7 min).

After the experiment, participants completed the short version of the Speech, Spatial, and Qualities of hearing Scale (SSQ; Gatehouse and Noble, 2004) and filled in the Edinburgh Handedness Inventory (EHI; Oldfield, 1971). On average each session lasted ~2.5 hours including EEG preparation.

### Behavioural analysis

All statistical analyses were performed in R Studio (version 1.2.1335, R Core Team, 2016, Vienna, Austria) running the R software package (version 3.6). Circular statistics were performed by using the package ‘circular’ in R (Agostinelli and Lund, 2013) and with code provided by Pewsey et al. (2013) (Circular Statistics in R).

We applied signal-detection theory and calculated visual target sensitivity (d’) for each of the 20 visual target times. Trials were included in the analysis if a button press occurred between 100 and 1000 ms after the Gabor grating was presented (similar to Henry and Obleser, 2012; Henry et al., 2014, 2017; Bauer et al., 2018). Trials in which button presses occurred outside of this response window as well as trials in which no responses were made were discarded (*M* = 57 trials, *SEM* = 14.8; 8.88 ± 2.30%). In addition, trials were excluded if a participant blinked around the presentation of the Gabor grating (−500 to 100 ms), as quantified via vertical electrooculogram (*M* = 9 trials, *SEM* = 2.9; 1.46 ± 0.46%; see section EEG data acquisition and pre-processing). On average, 574 trials (*SEM* = 16.5; 89.65 ± 2.58%) were retained for the analysis of visual target sensitivity per participant.

### Analysis of visual target sensitivity

We applied Signal Detection Theory in order to calculate visual target sensitivity. To apply the signal-detection framework to our behavioural data we assigned all 45-degree targets as the “Signal” condition and all 135-degree targets as the “Noise” condition (Macmillan and Creelman, 1991; Zhang et al., 2019). The hit rate was calculated as the proportion of all trials with a 45-degree target in which a correct 45-degree response was made (while incorrect responses in 45-degree target trials were classified as miss; see Figure 1C). The false alarm rate was calculated as the proportion of all trials with a 135-degree target in which an incorrect 45-degree response was made (while correct responses in 135-degree target trials were classified as correct rejection). For each of the 20 visual target times we calculated the sensitivity (d’) with the following equation:

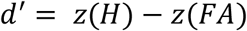

where *z*(H) refers to the z-score transformation of the hit rate and *z*(FA) refers to the z-score transformation of the false alarm rate (Macmillan and Creelman, 1991). We corrected for extreme proportions (0,1) by applying the log-linear rule, adding 0.5 to both the number of hits and the number of false alarms, and 1 to both the number of 45 and 135-degree trials (Hautus, 1995; Stanislaw, 1999; Ho et al., 2017). Visual target sensitivity was first calculated separately for each of the 20 visual target times. These were subsequently smoothed with an unweighted circular-moving average with a bin size of +/-2 (procedure similar to Henry and Obleser, 2012; Henry et al., 2017; Bauer et al., 2018).

### Analysis of cross-modal behavioural entrainment

Rhythmic modulations of behavioural performance in response to FM auditory stimulation have been previously observed in pure auditory tasks (Henry and Obleser, 2012; Henry et al., 2014, 2017; Bauer et al., 2018). To investigate the influence of the 3-Hz FM auditory stimulation on behavioural performance in the visual target identification task we derived two separate measures. First, we calculated a circular-linear correlation between visual target sensitivity and the phase of the 3-Hz FM stimulus at which the visual target was presented to test whether behavioural entrainment generalises to visual performance. Second, we fitted a single-cycle sine function to the behavioural data of each participant in turn. From this fit we extracted the phase angle corresponding to the best behavioural performance.

Circular-linear correlations between visual target sensitivity values and the 3 Hz FM stimulus phase were calculated separately for each participant to test whether visual target sensitivity was significantly modulated by the 3-Hz FM stimulus phase (Johnson-Wehrly-Mardia Correlation Coefficient; 1000 permutations). The circular-linear correlation can be interpreted as the degree to which behavioural performance is modulated by the FM phase and we will refer to the circular-linear correlations as cross-modal behavioural entrainment throughout the manuscript. To investigate the strength of these circular-linear correlations across participants, we performed permutation testing. For each participant, we formed a permutation distribution of circular-linear correlation coefficients by shuffling the correspondence between the 3Hz FM stimulus phase and visual target sensitivity values. Permuted data were smoothed using a circular-moving average (bin size +/-2) and circular-linear correlations were calculated on each of 1000 iterations. We obtained a z-score for the actual circular-linear correlation by

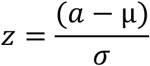

where *z* is the *z*-transformed observed data, *a* is the observed data (i.e., actual circular-linear correlation), and *μ* and *σ* are mean and standard deviation of the permutation distribution, respectively. The resulting z-scores were then tested against 0 using a one-sample *t*-test.

As mentioned above, we additionally fitted a single-cycle sine function to each participants’ behavioural data (visual target sensitivity) using the Levenberg Marquardt nonlinear least-squares algorithm implemented in the R package minpack.lm (Elzhov et al., 2015):

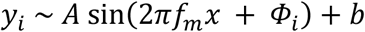

where *y_i_* are the observed behavioural data in relation to the time steps *x* ε [0, 0.33] in seconds for one cycle of the *f_m_* 3-Hz FM stimulation. *A, Φ_i_*, and b represent the amplitude, the phase lag of the sine fit, and the intercept respectively. For each participant, we obtained the best sine-fit function *g_i_*(*x*) = *A*sin(2π*f_m_x* + *Φ_i_*)+*b* by applying the nonlinear least-squares algorithm, allowing amplitude, phase lag, and intercept to vary (three degrees of freedom). From the best-fitting sine functions for each participant we calculated the phase angle corresponding to peak performance by estimating the local maximum of *g_i_* (*x*_max_ = arg max_*x*_ *g_i_*(*x*)). We obtained the phase angle in radians for peak performance by multiplying *x*_max_ by 2π *f_m_*, where *f_m_* = 3 Hz for each participant. To test whether there is a systematic relation between visual performance and auditory phase across participants, optimal stimulus phases were tested for uniformity, using the Rayleigh test.

### EEG data acquisition and pre-processing

EEG was acquired using Synamps amplifiers and Neuroscan acquisition software (Compumedics Neuroscan). We used a custom 62-channel setup with the following subset of electrodes of the international 10-10 system: FPz, AFz, AF3/4, AF7/8, Fz, F1/2, F3/4, F7/F8, FCz, FC1/2, FC3/4, FC5/6, FT7/8, Cz, C1/2, C3/4, C5/6, T7/8, CPz, CP1/2, CP3/4, CP5/6, TP7/8, Pz, P1/2, P5/6, P7/8, P9/10, POz, PO3/4, PO7/8, PO9/10, Oz, O1/2, Iz, I1/2. The left mastoid was used as the online reference and we included a right mastoid measurement to derive an average-mastoid reference offline. The ground electrode was placed at the left upper arm. Two bipolar electrode pairs were used to record electrooculography; one pair was placed above and below the left eye (vertical electrooculography) and another lateral of each eye (horizontal electrooculography). During acquisition, signals were low-pass filtered by an antialiasing filter (250 Hz cut-off), digitized at 1000 Hz, and stored for offline analysis.

EEG data were analysed in Matlab version 2017b (Mathworks), using a combination of EEGlab (version 14_1_2b used for pre-processing; Delorme and Makeig, 2004), Fieldtrip (version 20190419 used for analysis of inter-trial phase coherence; (Maris and Oostenveld, 2007), and custom scripts. For the ICA decomposition EEG raw data were first filtered offline between 1 and 40 Hz (finite impulse response filter, filter order high-pass filter: 800; filter order low-pass filter: 100), down-sampled to 500 Hz and EEG data between task blocks (i.e. during breaks) were pruned. Data were subsequently segmented into consecutive 1-s time intervals, and segments containing non-stereotypical artefacts - defined as epochs with a joint probability greater than 3 standard deviations from means of local (single-channel) and global (across channels) activity distributions - were rejected (pop_jointprob.m, locthresh & globthresh = 3). The remaining data were submitted to an independent component analysis (ICA) based on the extended Infomax (Bell and Sejnowski, 1995; Jung et al., 2000a, 2000b). The resulting unmixing weights were used to linearly decompose the original and down-sampled raw data and attenuate typical artefacts that reflected eye blinks, horizontal eye movements, heartbeat, and other sources of non-cerebral activity (components identified via visual inspection; *M* = 9.7, *SEM* = 0.5 components removed per data set). The code used for this pre-processing step is publicly available by (Stropahl et al., 2018). The ICA-corrected raw data were subsequently filtered between 0.5 and 40 Hz (finite impulse response filter, filter order high-pass filter: 1600; filter order low-pass filter: 100)

We applied a surface Laplacian transform (Perrin et al., 1989) to the EEG data as implemented by (Cohen, 2014). The surface Laplacian transform was applied to increase spatial resolution and to obtain a reference-free representation of the underlying current generators (Kayser and Tenke, 2015). The onset phase of the auditory stimulation varied on a trial-by-trial basis and could take one of four values: 0, pi/2, pi, and 3pi/2. In order to test for neural entrainment effects, EEG data were epoched relative to the rising phase of the auditory stimulation (phase 0 of the sine wave), so that the auditory stimulus phase was consistent across trials. Specifically, EEG data were epoched from −2000 to +4000 ms relative to the re-aligned auditory stimulation onset and baseline-corrected in the time window of −150 to 0 ms. As with the behavioural data, we only considered trials in which responses occurred within 100 to 1000 ms after visual target presentation. Trials with no button presses or responses outside of this response window were discarded. Further, we rejected all trials in which eye blinks occurred from −500 to +100 ms relative to visual target onset (*M* = 9 trials, *SEM* = 2.9; 1.46 ± 0.46%). Eye blinks were detected via vertical electrooculography prior to the ICA decomposition. In particular, trials on which the vertical EOG voltage surpassed ~200μV (approximately one-half of the maximum voltage evoked by a typical blink) were flagged and confirmed by visual inspection. Finally, epochs with an especially high variance were discarded (pop_jointprob.m, locthresh & globthresh = 5). On average 532 trials (*SEM* = 15.4; 83.15 ± 2.37%) were retained for the EEG analysis per participant.

### EEG entrainment and statistical analysis

To investigate effects of neural entrainment in response to the auditory stimulation we calculated inter-trial phase coherence (ITPC; Lachaux et al., 1999). To this end, a fast Fourier transform was calculated, including a Hanning taper and zero padding, for each trial and each channel using a fixed number of 6 cycles across frequencies. The resulting complex numbers were normalised by dividing each complex number by its magnitude. ITPC was then calculated as the absolute value of the mean normalised complex number across trials. ITPC values can take on values between 0 (no coherence) and 1 (perfect phase coherence). ITPC was calculated for frequencies ranging from 1 to 10 Hz in steps of 0.1 Hz and in time steps of 100 ms. In an initial step, we calculated ITPC across all trials. Subsequently the analysis was repeated to determine ITPC values separately for trials in which visual targets occurred during the first (~1000 - 1333 ms) or second cycle (~1333 - 1667 ms) of the visual target window (see Figure 1B). The motivation for breaking down the ITPC analysis was twofold. First, we wanted to ensure that the reported effects in visual electrodes were not contaminated by evoked responses due to the visual target. Second, we wanted to confirm that auditory and visual electrodes demonstrated distinct response patterns if the visual target occurred in the first or second cycle (see Figure 4).

Statistical analyses were performed separately for electrodes associated with auditory and visual brain activity. For auditory activity, four bilateral electrodes were chosen based on the 3-Hz ITPC topography (FC3, FC5, FC4, FC6, TP7, P7, TP8, P8). The electrodes chosen correspond well with the scalp distributions of auditory generators observed in previous studies that also applied a Laplacian transform (e.g. Jaeger et al., 2018). For visual activity, we chose to use the three central occipital electrodes *a priori* (O1, Oz, O2) to investigate potential entrainment of visual activity. We focused on these canonical visual electrodes because our visual targets were presented centrally, and because these electrodes were the most remote from any possible effect of entrainment on auditory brain activity. In addition to using a selected number of electrodes, we also calculated ITPC values for all electrodes and projected them on the electrode array to obtain a topographical visualisation of the ITPC patterns (see Figure 3A).

For statistical analyses, we extracted ITPC values for both auditory and visual electrodes at the 3-Hz stimulation frequency by calculating the average across the 2.5 to 3.5-Hz frequency window and the 500 to 1000 ms time window. The starting point of the time window was based on a previous study showing reliable auditory entrainment after 2 to 3 cycles of a 3-Hz FM stimulus (Bauer et al., 2018). The end point was chosen to avoid contamination by brain activity related to visual target onset or responding.

To compare the strength of peristimulus ITPC against baseline ITPC, paired sample *t*-tests were calculated, separately for the selected auditory and visual electrodes. The baseline ITPC was defined as the average 2.5-to 3.5-Hz ITPC in the −800 to −300 ms time window before stimulation onset. This segment was chosen as it does not overlap with the realignment of trials nor potential backward smearing cause by the sliding time window used for the ITPC analysis.

In addition, we investigated neural entrainment strength as a function of behavioural performance. We used the k-means algorithm by (Hartigan and Wong, 1979) as implemented in R (R Core Team, 2016) to cluster participants into three groups based on their cross-modal behavioural entrainment data (i.e. circular-linear correlations). K-means algorithm is an iterative algorithm that partitions the dataset into k-groups such that the sum of squares from points to the assigned cluster centre is minimised. We will refer to these three groups as high (*N* = 10), medium (*N* = 11), and low cross-modal behavioural entrainment group (*N* = 7). The number of participants in each group was not set a-priori but instead was defined by the clustering algorithm. After having found these groups based purely on the behavioural data, we performed a contrast analysis with the ITPC values as dependent measure to investigate whether there is a linear trend among the means of the three groups. The contrast analysis was followed up by a Spearman rank order correlation to test the relationship between behavioural and neural entrainment on a more continuous scale.

As phase coherence values are bounded between 0 and 1 and are therefore not normally distributed, ITPC values were arcsine-transformed before being submitted to statistical analysis (Studebaker, 1985). Throughout the manuscript, effect sizes are provided as Cohens *d* for *t*-tests. All statistical tests are two-tailed and the significance level was set to *p* < 0.05 for all tests.

## Results

### Auditory rhythmic stimulation modulates visual target sensitivity

Visual target sensitivity was not stable across visual target times, but systematically co-varied with the phase of the auditory stimulation. Based on our staircasing of the visual target contrast, the mean d’ calculated over all target times (target times depicted in Figure 1B) showed that participants performed the task better than chance, but far from ceiling (*M* = 1.57, *SEM* = 0.10). Critically, however, around the mean values, behavioural performance ebbed and flowed systematically according to the phase of the auditory 3-Hz FM stimulation. Figure 2A shows the behavioural data of all 28 participants in descending order from high to low cross-modal behavioural entrainment. Circular-linear correlations between the 3-Hz FM stimulus phase and the visual target sensitivity values were systematically higher than would be expected by chance (*t*-test against 0, two-tailed: *t*(27) = 8.628, *p* < 0.001, *d* = 1.630). Even at the level of individual participants, circular-linear correlations reached significance for 21 out of 28 participants (Figure 2A).

**Figure 2.**
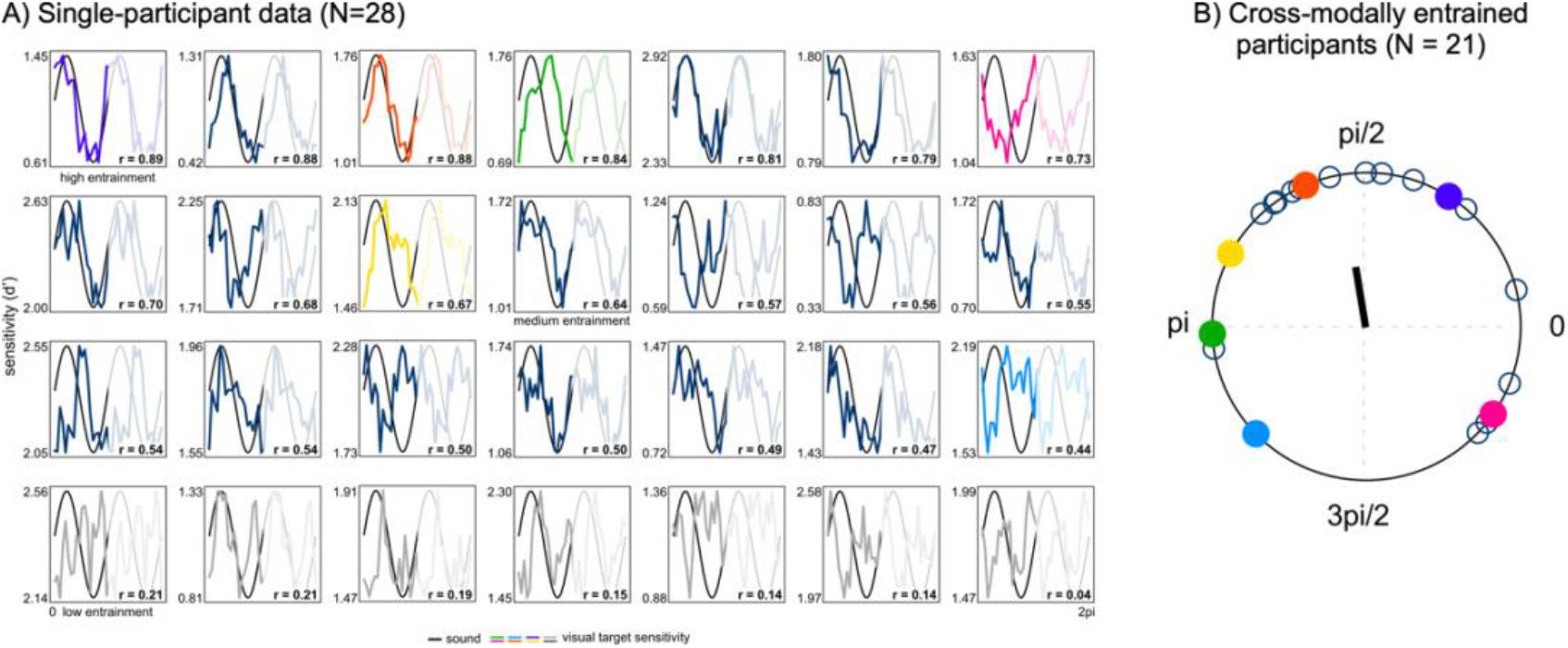
Rhythmic behavioural modulation of visual discrimination performance by auditory FM stimulation. A) Single-participant data for all 28 participants ordered in descending order from high to low cross-modal behavioural entrainment (i.e. circular-linear correlation; correlation value is shown in each plot). Plots show smoothed behavioural performance for all 20 visual target times (coloured and grey/dark blue lines) superimposed on a schematic of the sound (black line). The colour lines correspond to participants as shown in B. Two cycles were concatenated for better visualisation purposes (2^nd^ cycle is faded). Further, participants were divided into three groups: high (N = 10), medium (N = 11), and low cross-modal behavioural entrainment (N = 7) group. Groups were based on k-means clustering and used for the analysis of the neuro-behavioural relationship (see Figure 5). Participants in grey did not show a significant cross-modal behavioural entrainment (circular-linear correlation: *p* > 0.05). B) Phase angle distribution for *N* = 21 participants showing significant cross-modal behavioural entrainment. Data from seven participants were removed (grey lines in A). Colours correspond to the single-participant data as shown in A. The black arrow indicates the mean phase angle across participants.

While the observed coupling was highly robust, we also found considerable variability across participants in the *phase* of this cross-modal behavioural entrainment pattern – such that visual target sensitivity peaked at different phases of the 3-Hz FM signal in different participants (compare for example the green and the pink participant in Figure 2A). In fact, we found no support for a clear phase concentration of this behavioural entrainment when considering the full sample (*N* = 28, Rayleigh *z* = 1.778, *p* = 0.252; *θ* = 108.2°, *SEM* = 17.9°). However, when excluding the 7 participants for whom we did not observe significant cross-modal behavioural entrainment in the first place (circular-linear correlation *p*-values > 0.05; grey lines in Figure 2A), significant phase clustering became evident in the group (Figure 2B, *N* = 21, Rayleigh *z* = 3.212, *p* = 0.038; *θ* = 98.4°, *SEM* = 17.1°).

Thus, despite variability in the observed phase of entrainment, cross-modal behavioural entrainment of visual target perception by the 3-Hz FM stimulation was highly robust, and could be identified in the majority of participants.

### Auditory FM stimulation modulates activity in auditory electrodes

We focused our EEG analyses on the 21 participants who showed significant evidence for a behavioural modulation by the 3-Hz FM stimulation. As a main indicator for neural entrainment we calculated inter-trial phase coherence (ITPC; Lachaux et al., 1999). In line with previous studies showing neural entrainment in response to a FM stimulus (Henry and Obleser, 2012; Henry et al., 2014, 2017; Bauer et al., 2018), we also found clear evidence for entrainment of 3-Hz activity to the FM stimulation (Figure 3B). The 3-Hz topography for ITPC values averaged over the 2.5 - 3.5 Hz frequency and 500 - 1000 ms (Figure 3A) shows a clear bilateral modulation of auditory processing with a similar distribution as previous auditory studies that also used a Laplacian transform (e.g. SanMiguel et al., 2013; Kayser and Tenke, 2015; Jaeger et al., 2018). A paired-samples *t*-test confirmed a significant increase in ITPC in the designated auditory electrodes, relative to the pre-stimulation baseline (Figure 3B; *t*(20) = 4.676, *P* < 0.001, *d* = 1.02; *M*_baseline_ = 0.05, *SEM*_baseline_ = 0.001, *M*_stimulation_ = 0.09, *SEM*_stimulation_ = 0.01).

**Figure 3.**
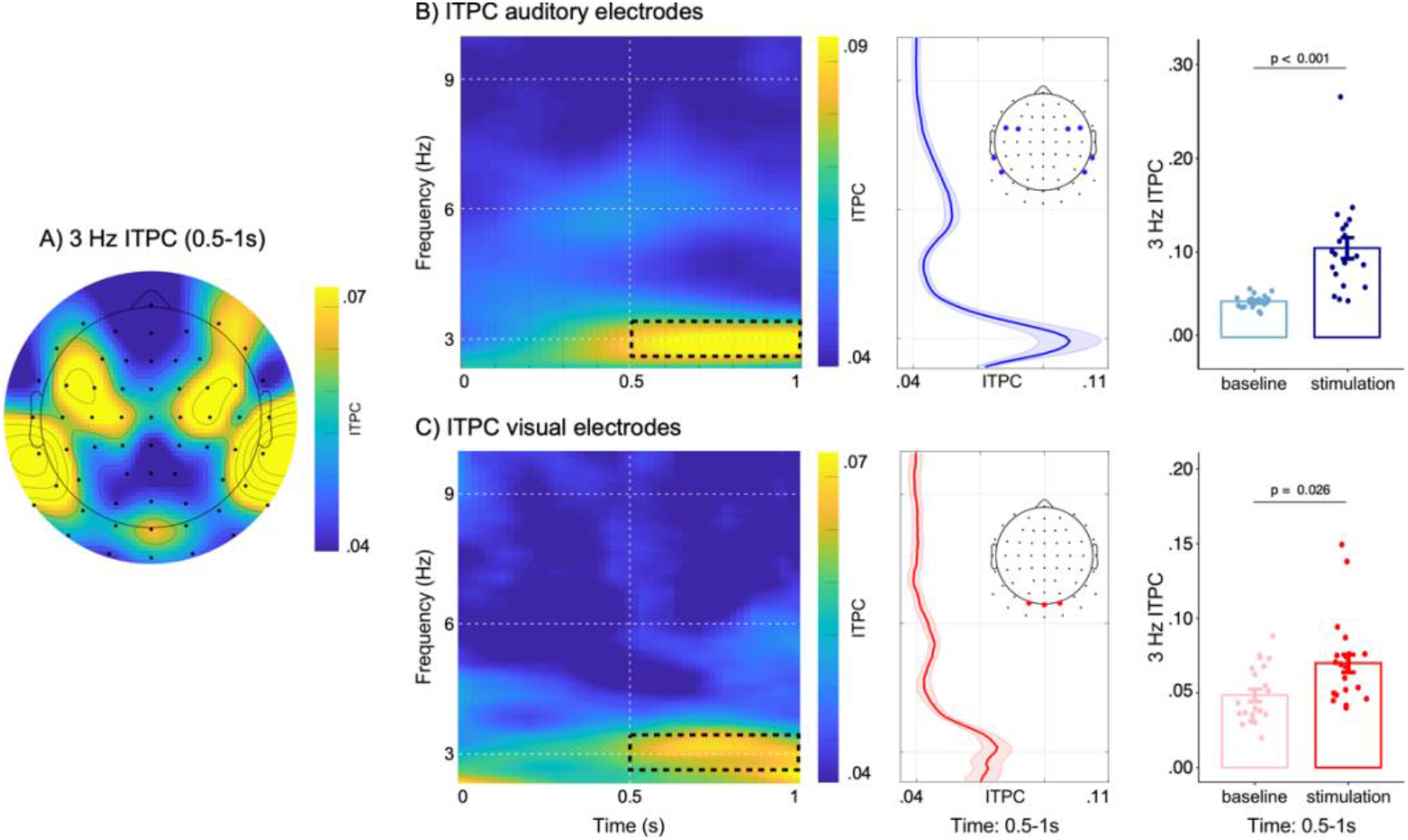
Entrainment of neural activity of auditory and visual EEG electrodes. Data for *N* = 21 participants with significant behavioural cross-modal entrainment A) 3-Hz topography, averaged for the 2.5 to 3.5 Hz frequency range and 500-1000 ms time period. A surface Laplacian transform was applied to help separate contributions from auditory and visual areas. B) ITPC in auditory electrodes. Left panel shows the time-frequency representation with a clear 3-Hz activation. Black box indicates the time (500 to 1000 ms) and frequency range (2.5 to 3.5 Hz) that was used for statistical analysis. Middle panel shows ITPC values collapsed over the time window of 500-1000 ms; shaded area depicts the standard error of the mean and the inset topography depicts the electrodes used for statistical analysis. Bar plots in the right panel depict extracted 3-Hz ITPC values (2.5 to 3.5 Hz) for both, the **stimulation period** (500 to 1000 ms; dark blue) and the pre-stimulus baseline period (−800 to −300 ms; light blue). C) ITPC in visual electrodes. Time-frequency representation of ITPC values averaged across visual electrodes shows an increase in phase coherence at the 3-Hz stimulation frequency (black box indicates time and frequency range for statistical analysis). ITPC averaged over time is shown in the middle panel; shaded area shows the standard error of the mean. Right panel shows bar plots of the extracted 3-Hz ITPC values for stimulation and baseline periods.

### Auditory FM stimulation modulates activity in visual electrodes

In addition to auditory activations, the 3-Hz topography shown in Figure 3A also reveals an increase in ITPC over posterior electrodes. Such a pattern has not previously been observed in purely auditory (i.e., unimodal) studies (e.g. SanMiguel et al., 2013; Kayser and Tenke, 2015; Jaeger et al., 2018) and suggests entrainment of visual processing. A paired *t*-test in canonical and a-priori-defined visual electrodes (O1, Oz, O2) confirmed a significant increase in ITPC relative to the pre-stimulation baseline (Figure 3C; *t*(20) = 2.399, *p* = 0.0263, *d* = 0.52; *M*_baseline_ = 0.05, *SEM*_baseline_ = 0.003, *M*_stimulation_ = 0.07, *SEM*_stimulation_ = 0.005). These data thus suggest that in our cross-modal task, the 3-Hz FM stimulation did not only entrain neural activity in auditory electrodes, but also entrained oscillatory visual activity in posterior electrodes in the absence of any continuous visual stimulation and in a fashion consistent with the cyclical pattern of visual target sensitivity in behaviour that we reported in Figure 2A.

In a supplementary analysis, we could demonstrate clear 3-Hz ITPC in visual electrodes even for trials in which the visual target occurred relatively late in the auditory stimulation (Figure 4B) – making it unlikely that these reported effects in visual electrodes were dependent on the visual stimulation. Moreover, this analysis confirmed distinct response patterns associated with visual target processing between the selected visual and auditory electrodes. While visual target responses were evident in visual electrodes (Figure 4B) this was much less so in auditory electrodes (Figure 4A), thereby increasing our confidence in the separation of neural activity between the selected visual and auditory electrodes.

**Figure 4.**
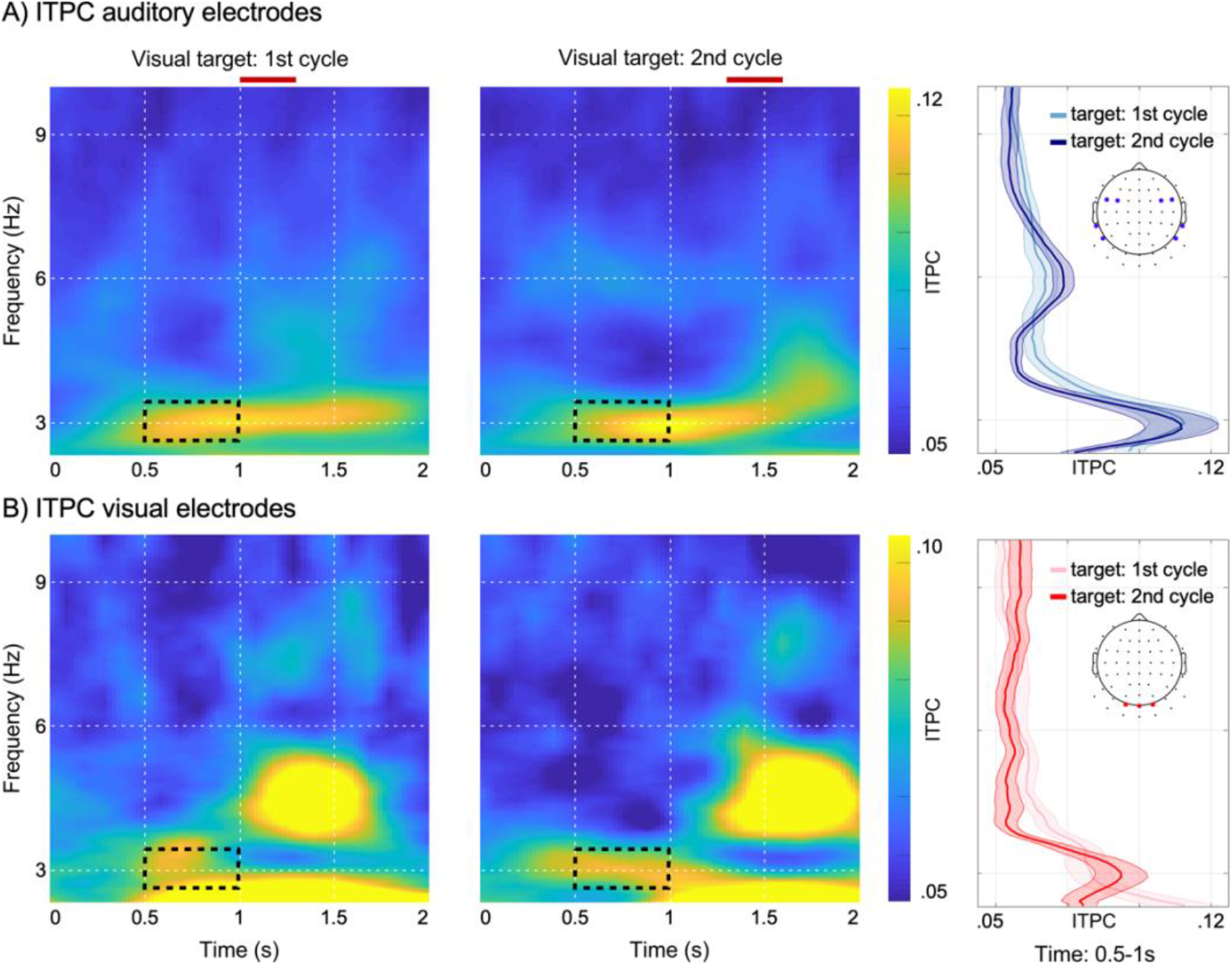
Entrainment of neural activity in auditory and visual EEG electrodes separated by target occurrence. ITPC analysis separately for visual targets occurring during first (~1000 to 1333 ms) or second cycle (~1333 to 1667 ms) of the visual target window (N = 21 participants are shown; see Figure 1B for visual target window). Visual target windows are indicated via red bars on top of the time-frequency representations. A) ITPC in auditory electrodes: left and middle panel show time-frequency representations for the first and second cycle of the visual target window. Right panel shows ITPC values collapsed over time (500 - 1000 ms) separately for targets occurring during the first (light blue) or second cycle (dark blue); shaded areas depict the standard error of the mean and the inset topography depicts the electrodes used for plotting. B) ITPC in visual electrodes: left and right panels show time-frequency representation of ITPC values averaged across visual electrodes for trials in which visual targets were presented in the first (left) and second (middle) cycle. ITPC averaged over the time window of 500 - 1000 ms is shown in the middle panel separately for targets occurring in the first (light red) and second cycle (dark red); shaded areas show the standard error of the mean and inset topography indicates electrodes used for plotting.

### Neural entrainment in visual electrodes is related to cyclic modulation of visual perception

Finally, we found that the degree of neural entrainment in visual electrodes was related to the degree to which visual target sensitivity fluctuated with the 3-Hz FM stimulation (Figure 5A,B).

**Figure 5.**
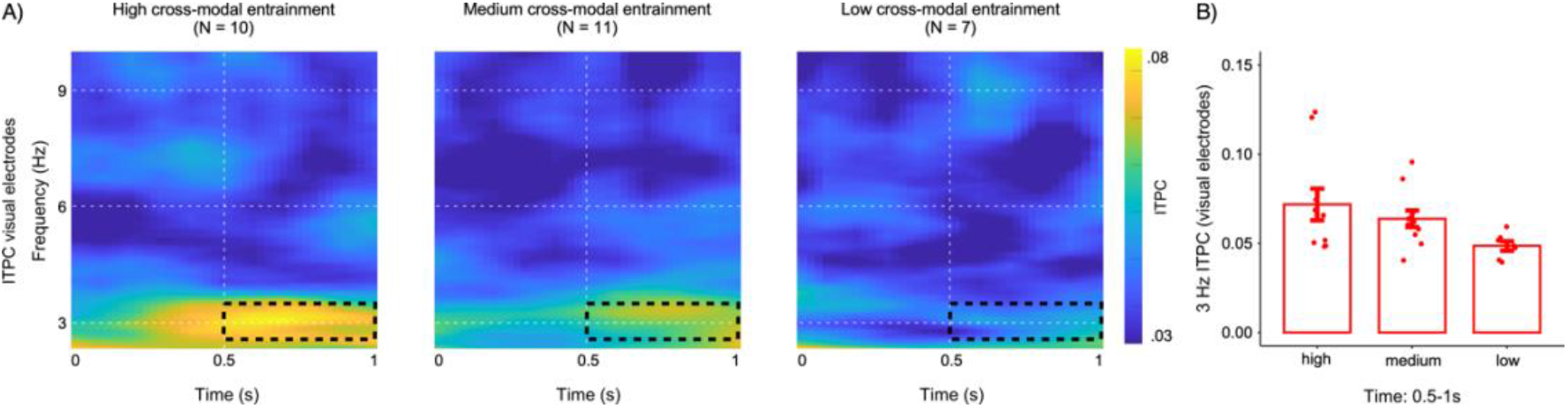
Neural entrainment in visual electrodes predicts the degree of cross-modal entrainment of visual perception. Data are shown for all participants (*N* = 28). A) Time-frequency maps for visual electrodes shown separately for the high, medium, and low cross-modal behavioural entrainment group. Dashed black boxes indicate the time-frequency range used for statistical analysis (2.5 - 3.5 Hz and 500 - 1000 ms). B) Red bar plots depict 3-Hz ITPC values for visual electrodes for each group.

A k-means algorithm (Hartigan and Wong, 1979) partitioned individuals who had a low (*N* = 7; cluster centre = 0.154), medium (*N* = 11, cluster centre = 0.528), or high (*N* = 10, cluster centre = 0.787) cross-modal behavioural entrainment in a purely data-driven manner. While this sorting was performed exclusively on the pattern of behavioural data, we found striking differences in the pattern of visual ITPC between the groups (Figure 5A). For example, while neural entrainment in visual electrodes was prominent in the group that also showed clear behavioural entrainment (Figure 5A, left panel), we found virtually no entrainment in visual activity in the group that also showed no clear behavioural modulation (Figure 5A, right panel; note that this group also corresponds to the 7 participants for which we could not establish a significant effect in behaviour). A contrast analysis comparing ITPC in visual electrodes across the three groups (low, medium, high) showed a significant linear trend among the means of the three groups (*F*(1,25) = 5.658, *p* = 0.025), whereby neural entrainment in visual electrodes was largest in the participant group that also showed the largest cross-modal behavioural entrainment. Arguing for the functional relevance of visual entrainment, this linear trend among the three groups was corroborated by a significant positive correlation between ITPC values in visual electrodes and cross-modal behavioural entrainment (*rho*(27) = 0.39, *p* = 0.039) across participants.

Strikingly, when running the same analyses for the neural entrainment activity in *auditory* electrodes we found no linear trend among the means of the three groups (*F*(1,25) = 1.954, *p* = 0.174), nor a significant correlation (*rho*(27) = 0.22, *p* = 0.27).

## Discussion

The current study provides a strong validation of cross-modal entrainment as a useful means for organising the perception of multisensory stimulation in the natural environment (Lakatos et al., 2019). Our results yielded clear behavioural evidence for cross-modal auditory-to-visual entrainment in response to a continuous FM stimulation. Further, our findings show that visual cortical activity was modulated by the 3-Hz FM stimulation in the absence of any transient changes in stimulus energy or salience in the auditory stream and without a concomitant continuous visual stimulus. Moreover, we report that the degree of cross-modal auditory-to-visual neural entrainment is related to the degree to which visual target sensitivity fluctuated with the 3-Hz FM stimulus. Together, we could demonstrate a cross-modal influence of auditory rhythmic stimulation on visual neural activity that is functionally relevant for behavioural performance.

Our study provides clear evidence for cross-modal entrainment through the careful choice of stimulus materials that remove likely alternative explanations to any rhythmic modulation of behaviour or brain activity. Building on previous work using FM stimulation in purely auditory (i.e. unimodal) settings (Henry and Obleser, 2012; Henry et al., 2014, 2017; Bauer et al., 2018), the constant amplitude of the FM stimulation allowed us to systematically present visual targets in relation to the phase of the auditory stimulation and thereby investigate a behavioural modulation profile that is not tied to any salient onset detections, nor potential masking effects introduced by abrupt on- or offsets.

Nevertheless, the nature of the FM stimulus leaves open some alternative explanations that will require further investigation. While we used a FM-stimulation to diminish the perception of onset and offset effects at the cortical level, it might not necessarily be the case that the current findings are due to a genuine entrainment of ongoing brain oscillations. For example, participants might have perceptually “parsed” the sounds into high- and low-frequency periods resulting in evoked perceptual onsets (though the heterogeneity in the preferred “phase” of behavioural entrainment argues against any obvious “anchoring point” in the FM-sound). It is worth considering, of course, that modulation of rhythmic neural activity in accordance to changes in the properties of sensory stimuli is a natural process of information processing and need not always be discounted as artefactual. The important question in our study was whether these modulations in auditory rhythmic activity had any consequence for visual performance and visual neural activity.

The observed modulations of visual performance go beyond earlier work using either uni-modal or cross-modal rhythmic stimulation. Our use of signal detection theory enabled us to demonstrate effects of cross-modal entrainment on perceptual sensitivity, while controlling for potential changes in response bias. The results therefore expand on findings from previous studies that have documented cyclic modulation on detection rates and reaction times but without controlling for possible changes in criterion shifts, as reported, for example, during uni-modal auditory (Henry and Obleser, 2012; Henry et al., 2014; Bauer et al., 2018) or visual (Busch et al., 2009; Mathewson et al., 2009; Busch and VanRullen, 2010; Samaha et al., 2015) rhythmic stimulation. Further, studies using cross-modal rhythmic stimulation have mainly tested on-vs. off-beat target presentations showing enhanced visual performance for targets occurring on beat with a preceding auditory rhythm (Miller et al., 2013; Escoffier et al., 2015; Simon and Wallace, 2017; Barnhart et al., 2018). Here, we were able to construct a behavioural modulation profile for each individual participant. Moreover, while previous studies investigating cross-modal entrainment focused either on behavioural (Miller et al., 2013; Barnhart et al., 2018) or neural activity (Lakatos et al., 2008; Escoffier et al., 2015) we were able to link cross-modal neural entrainment effects with behavioural performance. This link between neural entrainment of visual activity and behavioural performance suggests a functional relevance of the cross-modally entrained activity for behaviour.

The observed link between behavioural and neural entrainment was driven by substantial differences in the degree of entrainment across participants. One explanation might be that some participants were better able to focus solely on the visual task, and hence to ignore the auditory stimulation. This would lead to little cross-modal entrainment of neural activity and behaviour. Another possibility is that cross-modal entrainment varies between participants for other reasons (such as hardwired anatomical connectivity), independent of potential strategic factors. In addition, we also observed heterogeneity in the preferred phase angle across participants. Previous studies using pure auditory stimulation observed similar variability across participants (Henry and Obleser, 2012, 2013; Henry et al., 2014; Bauer et al., 2018). A possible reason for this heterogeneity is that some participants may be faster in adapting to the 3-Hz FM stimulation than others and may have different neural lags that could account for the variability (Besle et al., 2011; Henry and Obleser, 2012; Bauer et al., 2018). Further, differences in perceptual parsing of the 3-Hz FM sound might explain some variability among the participants as well as individual difference in intrinsic brain rhythms (Kösem et al., 2014).

The pattern of rhythmic modulation of EEG activity in canonical visual electrodes suggests rhythmic entrainment of visual cortical processing. Whereas one may worry that the observed visual effect merely reflects volume conduction, we have reasons to believe that the observed visual entrainment reflected more than just volume conduction. First, the increase in visual ITPC as seen in the current study, has not been reported in purely auditory studies using a similar pre-processing pipeline and a Laplacian surface filter (SanMiguel et al., 2013; Kayser and Tenke, 2015; Jaeger et al., 2018). Second, our observed neuro-behavioural relationship was only significant for the ITPC in the visual electrodes, but not the auditory electrodes. If this reflected volume conduction, this correlation should have been clearer for the directly driven auditory electrodes where the neural entrainment was (not surprisingly) much larger. Third, our cycle-by-cycle analysis showed distinct activation patterns for auditory and visual electrodes.

There are multiple possible routes for auditory-to-visual entrainment. Cross-modal auditory-to-visual influences in neural activity could occur through a direct influence between auditory and visual cortices as suggested by animal studies (Kayser et al., 2008; Lakatos et al., 2009; Perrodin et al., 2015). Alternatively, or additionally, information could be transferred indirectly through higher-order multisensory regions (i.e. superior temporal sulcus, intraparietal sulcus, and prefrontal cortex; Ghazanfar and Schroeder, 2006; Driver and Noesselt, 2008; Van Atteveldt et al., 2014), or subcortical regions (Cappe et al., 2007; Hackett et al., 2007; Lakatos et al., 2007). Furthermore, there is growing evidence that neural oscillations can be modulated by top-down attention-related processes in multisensory settings (Keil et al., 2016; Macaluso et al., 2016; Auksztulewicz et al., 2017). These various potential routes for auditory-to-visual influences are mutually compatible. Temporal delay in information transfer in direct or indirect connections and variability in the focus and strength of factors linked to expectation and attention may also all contribute to the observed variability in preferred phase angles across participants. The current study design cannot address whether the observed cross-modal entrainment effects arise due to a bottom-up auditory-to-visual drive, indirect or re-entrant connections, top-down attention-related factors, or interactions among these sources. Future research on cross-modal entrainment can shed light on the underlying mechanisms, for example by directly manipulating attention or temporal expectations of visual target occurrence.

Taken together, we have shown that a continuous auditory rhythm acts as a pacing signal by which neural oscillations can be entrained cross-modally and thereby guide visual behavioural performance ensuring that incoming visual stimuli are efficiently processed. An outstanding question is whether this tracking of environmental rhythms is special for the auditory domain, or whether, for example, a continuous visual rhythm proves equally effective to induce rhythmic modulations in the auditory domain. Further, auditory rhythms to investigate cross-modal auditory-to-visual influences were so far restricted to the delta and theta frequency range, which conform with the range in which endogenous brain oscillations operate (Lakatos et al., 2008). Whether the observed cross-modal rhythmic modulations of visual perception can operate outside of this frequency range needs to be further investigated (see also Zalta et al., 2020). Even so, the current study provides an important addition to the cross-modal entrainment hypothesis by showing cross-modal auditory-to-visual influences on brain and behaviour – in the absence of salient stimulus on- or offsets – and with behavioural consequences for perception.

## Acknowledgments

This research was funded by the German Research Foundation (DFG: MA 8554/1-1) to A.-K.R.B., a Marie Skłodowska-Curie Fellowship from the European Commission (ACCESS2WM) and an ERC Starting Grant from the European Research Council (MEMTICIPATION, 850636) to F.v.E., and a Wellcome Trust Senior Investigator Award (ACN) (104571/Z/14/Z) and a James S. McDonnell Foundation Understanding Human Cognition Collaborative Award (220020448) to A.C.N., and the NIHR Oxford Health Biomedical Research Centre. The Wellcome Centre for Integrative Neuroimaging is supported by core funding from the Wellcome Trust (203139/Z/16/Z). The funders had no role in study design, data collection and analysis, decision to publish or preparation of the manuscript. We also wish to thank Carsten Matke for helpful discussions and comments.

## Author contributions

A.-K.R.B. and A.C.N. designed research; A.-K.R.B. performed research; A.-K.R.B., F.v.E. and A.J.Q. analysed data; A.-K.R.B., F.v.E., A.J.Q., and A.C.N. wrote the paper.

